# Differences in signalling, trafficking and glucoregulatory properties of glucagon-like peptide-1 receptor agonists exendin-4 and lixisenatide

**DOI:** 10.1101/803833

**Authors:** Philip Pickford, Maria Lucey, Zijian Fang, Stavroula Bitsi, Johannes Broichhagen, David J. Hodson, James Minnion, Guy A Rutter, Stephen R Bloom, Alejandra Tomas, Ben Jones

**Author notes:** Corresponding authors: Ben Jones and Alejandra Tomas. Contributed equally (Philip Pickford, Maria Lucey, Alejandra Tomas, Ben Jones).

## Abstract

**Background and purpose:** Amino acid substitutions at the N-termini of glucagon-like peptide-1 receptor agonist (GLP-1RA) peptides result in distinct patterns of intracellular signalling, sub-cellular trafficking and efficacy *in vivo*. Here we aimed to determine whether sequence differences at the ligand C-termini of clinically approved GLP-1RAs exendin-4 and lixisenatide lead to similar phenomena. We also sought to establish the impact of the C-terminus on signal bias resulting from modifications elsewhere in the molecule.

**Experimental approach:** Exendin-4, lixisenatide, and N-terminally substituted analogues with biased signalling characteristics were compared across a range of *in vitro* trafficking and signalling assays in different cell types. Fluorescent ligands and new time-resolved FRET approaches were developed to study agonist behaviours at the cellular and sub-cellular level. Anti-hyperglycaemic and anorectic effects of each parent ligand, and their biased derivatives, were assessed in mice.

**Key results:** Lixisenatide and exendin-4 showed equal binding affinity, but lixisenatide was 5-fold less potent for cAMP signalling. Both peptides were rapidly endocytosed, but the GLP-1R recycled more slowly to the plasma membrane after lixisenatide treatment. These combined deficits resulted in reduced maximal sustained insulin secretion and reduced anti-hyperglycaemic and anorectic effects in mice. N-terminal substitutions to both ligands had favourable effects on their pharmacology, resulting in improved insulin release and lowering of blood glucose.

**Conclusion and implications:** Changes to the C-terminus of exendin-4 affect signalling potency and GLP-1R trafficking via mechanisms unrelated to GLP-1R occupancy. These differences were associated with changes in their ability to control blood glucose and therefore may be therapeutically relevant.

## 1 Introduction

The glucagon-like peptide-1 receptor (GLP-1R) is a well-established pharmacological target for the treatment of both type 2 diabetes (T2D) and obesity due to its beneficial effects on weight loss and pancreatic beta cell function (Andersen et al., 2018). The main endogenous ligand for GLP-1R, the 29 amino acid peptide GLP-1(7-36)NH_2_, is highly susceptible to degradation by proteolytic enzymes that rapidly destroy it in the circulation, making it unsuitable as a therapeutic agent (Deacon et al., 1998). Therefore, a number of synthetic GLP-1R agonists (GLP-1RAs) with longer circulatory half-lives have been developed and subsequently approved for human use (Graaf et al., 2016). One example is the GLP-1 homologue peptide exendin-4 (Eng et al., 1992), in clinical use for T2D as exenatide. This molecule features an extended, proline-rich C-terminal extension (sequence GAPPPS-NH_2_) which is absent in GLP-1 itself. The precise role of this feature is not clear, but various possibilities have been suggested, including stabilisation of the peptide helical structure (Neidigh et al., 2001), facilitation of inter-protomer coupling within receptor oligomers (Koole et al., 2017), and protection against enzymatic degradation (Lee et al., 2018). A further approved T2D GLP-1 mimetic peptide, lixisenatide, shares a common first 37 amino acids with exendin-4, including most of the GAPPPS sequence, but includes an additional six lysine residues at the C-terminus prior to the terminal amidation (Andersen et al., 2018). Due to putative importance of the exendin-4 C-terminus, it is conceivable that the lixisenatide-specific changes could affect its pharmacology.

Recent work from our laboratory and others have highlighted how GLP-1R signal bias and related membrane trafficking effects regulate insulin release from beta cells (Zhang et al., 2015; Buenaventura et al., 2018; Jones et al., 2018b). Following agonist binding, the GLP-1R is rapidly endocytosed, and whilst active G protein coupled receptors (GPCRs) can continue to generate intracellular signals within the endosomal compartment (Eichel and Zastrow, 2018), the availability of surface GLP-1Rs to extracellular ligand appears to be an important determinant of sustained insulinotropic efficacy in a pharmacological setting (Jones et al., 2018b). The GLP-1R ligand N-terminus interacts with the receptor core to instigate conformational rearrangements needed for stable engagement with intracellular signalling and trafficking effectors, while its C-terminus facilitates this process by establishing the correct orientation of the peptide through interactions with the receptor extracellular domain (ECD) (Graaf et al., 2016). Suggesting that the C-terminal sequence differences between exendin-4 and lixisenatide might impact on these cellular processes, a limited evaluation in our earlier study implied that lixisenatide displays reduced signalling potency and insulinotropism compared to exendin-4 (Jones et al., 2018b).

In the present study, we extended our earlier evaluation to include formal comparison of bias between cAMP signalling and endocytosis in different cell types, measurements of post-endocytic targeting to recycling and degradative pathways aided by a novel cleavable time-resolved FRET probe, and assessment of the impact of these changes on exendin-4 *versus* lixisenatide metabolic responses *in vivo*.

## 2 Methods

### 2.1 Materials

All peptides and fluorescent peptide conjugates were obtained from WuXi AppTec (Wuhan) Co. Ltd. SNAP-Lumi4-Tb, BG-SS-Lumi4-Tb, and homogenous time-resolved fluorescence (HTRF) reagents for cAMP (cAMP Dynamic 2) and insulin (Insulin High Range Assay) measurement were obtained from Cisbio. SNAP-Surface probes were obtained from New England Biolabs. NanoGlo live cell reagents were obtained from Promega. Exendin-4 fluorescent EIA kits were obtained from Phoenix Pharmaceuticals. All other standard laboratory chemicals or culture reagents were obtained from Sigma or Thermo Fisher unless otherwise specified. The sources of all plasmids are indicated in the relevant sections below.

### 2.2 Cell culture

HEK293 cells (RRID: CVCL_0045) stably expressing human SNAP-GLP-1R (“HEK293-SNAP-GLP-1R”) were generated by G418 selection and maintained in DMEM with 10% FBS, 1% penicillin/streptomycin and G418 (1 mg/ml). HEK293T (RRID: CVCL_0063) cells were maintained similarly but without G418. Monoclonal CHO-K1 cells stably expressing human SNAP-GLP-1R (“CHO-K1-SNAP-GLP-1R”) were generated by G418 selection, with single clones obtained by FACS and subsequently expanded; cells were maintained in DMEM with 10% DBS, 1% non-essential amino acids, 20 mM HEPES, 1% penicillin/streptomycin and 1 mg/ml G418. Wildtype INS-1 832/3 cells (Hohmeier et al., 2000) (a gift from Prof. Christopher Newgard, Duke University) were maintained in RPMI-1640 with 11 mM glucose, 10 mM HEPES, 2 mM glutamine, 1 mM sodium pyruvate, 50 µM β-mercaptoethanol, 10% FBS, and 1% penicillin/streptomycin. INS-1 832/3 cells lacking endogenous GLP-1R after deletion by CRISPR/Cas9 (Naylor et al., 2016) (a gift from Dr Jacqueline Naylor, MedImmune) were used to generate a polyclonal population expressing human SNAP-GLP-1R by G418 selection, and were maintained as for wild-type INS-1 832/3 cells, with the addition of G418 (1 mg/ml). MIN6B1 cells (Lilla et al., 2003) (a gift from Prof Philippe Halban, University of Geneva) were maintained in DMEM with 15% FBS, 50 µM β-mercaptoethanol and 1% penicillin/streptomycin. Transfections were performed using Lipofectamine 2000 according to the manufacturer’s instructions.

### 2.3 TR-FRET surface receptor binding assays

HEK293-SNAP-GLP-1R cells were labelled in suspension with SNAP-Lumi4-Tb (40 nM) for 1 hour at room temperature in complete medium. After washing and resuspension in HBSS containing 0.1% BSA and metabolic inhibitors (20 mmol/L 2-deoxygucose and 10 mmol/L NaN_3_) to prevent GLP-1R internalisation (Widmann et al., 1995), binding experiments were performed using FITC-conjugated ligands as described below, with TR-FRET measured in a Flexstation 3 plate reader (Molecular Devices) using the following settings: λ_ex_ = 335 nm, λ_em_ = 520 and 620 nm, delay 50 μs, integration time 400 μs. Binding was quantified as the ratio of fluorescent signal at 520 nm to that at 620 nm, after subtraction of ratio obtained in the absence of FITC-ligands. <u>Saturation binding experiments with FITC-ligands</u>: cells were treated with FITC-ligands over a range of concentrations for 24 hours at 4°C before measurement. Equilibrium binding constants were calculated using the “one site – specific binding” algorithm in Prism 8 (GraphPad Software). <u>Competition binding experiments at equilibrium</u>: cells were treated with a fixed concentration (10 nM) of exendin(9-39)-FITC in competition with a range of concentrations of unlabelled exendin-4 or lixisenatide for 24 hours at 4°C before measurement. Binding constants were calculated using the “one site – fit K_i_” algorithm in Prism 8, using the equilibrium dissociation constant for exendin(9-39)-FITC measured in the same experiment by saturation binding. <u>Competition kinetic binding experiments</u>: TR-FRET signals were measured at regular intervals before and after addition of different concentrations of exendin(9-39)-FITC, or different concentrations of unlabelled agonist in combination with a fixed concentration (10 nM) of exendin(9-39)-FITC at 37°C. Rate constants for association and dissociation of the unlabelled ligands were calculated using the “kinetics of competition binding” algorithm in Prism 8, using values for exendin(9-39)-FITC determined in the same experiment.

### 2.4 Cyclic AMP assays by HTRF

Cells were stimulated with agonist for the indicated period in their respective growth mediums but without FBS. Assays were performed at 37°C without phosphodiesterase inhibitors, except for with INS-1 832/3 and MIN6B1 cells, where 3-isobutyl-1-methylxanthine (IBMX) was added at 500 µM. At the end of the incubation period, cells were lysed, and cAMP determined by HTRF (cAMP Dynamic 2 kit, Cisbio) using a Spectramax i3x plate reader (Molecular Devices).

### 2.5 Dynamic signalling measurements via FRET biosensors for cAMP and PKA

HEK293T cells were transiently transfected with the FRET-based cAMP sensor ^T^Epac^VV^ (a kind gift from Prof Kees Jalink, Netherlands Cancer Institute) (Klarenbeek et al., 2011) using Lipofectamine 2000. CHO-K1-SNAP-GLP-1R cells stably expressing the AKAR4-NES (Herbst et al., 2011) biosensor, a kind gift from Prof Jin Zhang (Addgene plasmid 64727), were used to measure cytoplasmic PKA activation. Cells were suspended in HBSS, placed into black, clear-bottomed plates, and FRET was measured before and after agonist addition in a Flexstation 3 plate reader at 37°C using the following settings: λ_ex_= 440 nm, λ_em_ = 485 and 535 nm. FRET was quantified as the ratio of fluorescent signal at 535 nm to that at 485 nm after subtraction of background signal at each wavelength. Dynamic FRET changes were expressed relative to individual well baseline to improve precision.

### 2.6 Measurement of mini-G and β-arrestin recruitment using NanoBiT complementation

The plasmids for mini-G_s_, −G_i_ and −G_q_, each tagged at the N-terminus with the LgBiT tag (Wan et al., 2018), were a kind gift from Prof Nevin Lambert, Medical College of Georgia. The plasmid for β-arrestin-2 fused at the N-terminus to LgBiT was obtained from Promega (plasmid no. CS1603B118); this configuration was used due to previous success with another class B GPCR (Shintani et al., 2018). The SmBiT tag was cloned in frame at the C-terminus of the GLP-1R by substitution of the Tango sequence on a FLAG-tagged GLP-1R-Tango expression vector (Kroeze et al., 2015), a gift from Dr Bryan Roth, University of North Carolina (Addgene plasmid # 66295). HEK293T cells in 12 well plates were co-transfected for 24 hours with Lipofectamine 2000 using the following quantities of DNA: 0.5 µg each of GLP-1R-SmBiT and LgBiT-mini-G, or 0.05 µg each of GLP-1R-SmBiT and LgBiT-mini-G with 0.9 µg empty vector DNA (pcDNA3.1). Cells were resuspended in Nano-Glo dilution buffer and fumarazine (1:20) (Promega) and seeded in 96-well half area white plates. Baseline luminescence was measured over 5 minutes using a Flexstation 3 plate reader at 37°C before addition of ligand or vehicle. Luminescent signal was then serially monitored over 30 minutes, and responses were normalised to average baseline.

### 2.7 Measurement of receptor internalisation by DERET

This assay was adapted from previous descriptions (Roed et al., 2014; Jones et al., 2018b). HEK293-SNAP-GLP-1R cells were labelled in suspension with SNAP-Lumi4-Tb (40 nM) for 1 hour at room temperature in complete medium. After washing and resuspension in 24 µM fluorescein solution in HBSS, TR-FRET was monitored before and after agonist addition in a Flexstation 3 plate reader at 37°C using the following settings: λ_ex_ = 335 nm, λ_em_ = 520 and 620 nm, delay 400 μs, integration time 1500 μs. TR-FRET was quantified as the ratio of fluorescent signal at 520 nm to that at 620 nm after subtraction of background signal at each wavelength (simultaneously recorded from wells containing 24 µM fluorescein in HBSS but no labelled cells).

### 2.8 Measurement of GLP-1R recycling by TR-FRET

CHO-K1-SNAP-GLP-1R cells adhered in white 96-well half-area tissue culture-treated plates were labelled with BG-SS-Lumi4-Tb (40 nM unless indicated otherwise) for 30 minutes at 37°C. BG-SS-Lumi4-Tb is a cleavable SNAP-tag probe that allows release of the lanthanide moiety following reduction of its disulfide bond when exposed to reducing agents. After washing, BG-SS-Lumi4-Tb labelled cells were treated with agonist for 30 minutes at 37°C to induce GLP-1R internalisation, placed on ice to arrest further trafficking, and residual surface GLP-1R was de-labelled by cleavage of the lanthanide using the cell-impermeant reducing agent 2-mercaptoethane sulfonate (Mesna, 100 mM in TNE buffer, pH 8.6) for 5 minutes. After further washing, exendin(9-39)-FITC, added as a non-internalising FRET-acceptor for the GLP-1R recycling to the cell surface, was added at 10 nM in HBSS, and TR-FRET signals were sequentially monitored at 37°C as above. Signal was expressed relative to the baseline TR-FRET ratio established from the first three readings.

### 2.9 Endosomal FITC-ligand binding assay

HEK293-SNAP-GLP-1R cells were labelled in suspension with SNAP-Lumi4-Tb (40 nM) for 1 hour at room temperature in complete medium. Cells were then washed and resuspended in complete medium containing 100 nM exendin-4-FITC, lixisenatide-FITC, or no ligand, which was allowed to internalise over 30 minutes at 37°C. Cells were then placed on ice, washed with cold HBSS, and incubated for 10 minutes in cold acetic acid + 150 mM NaCl buffer, pH 2.9, to strip surface ligand. After a final wash, cells were resuspended in HBSS and returned to 37°C. TR-FRET signal was measured over 30 minutes in a Flexstation 3 plate reader using the same settings as for binding experiments.

### 2.10 TR-FRET measurement of FITC-ligand endocytosis

HEK293-SNAP-GLP-1R cells were labelled in suspension with SNAP-Lumi4-Tb (40 nM) for 1 hour at room temperature in complete medium. After washing and resuspension in HBSS containing 0.1% BSA, cells were plated at 4°C to arrest endocytosis, and pre-chilled FITC-ligands were allowed to bind for 3 hours. The plate was then moved to the plate reader at 37°C to initiate endocytosis, and TR-FRET was measured at regular intervals using the same settings as for binding experiments. Ligand uptake was quantified by monitoring the change in TR-FRET ratio over time relative to the baseline value from the average of the first two recordings.

### 2.11 Microscopes

<u>Confocal microscopy</u>: fixed, mounted coverslips were imaged with a Zeiss LSM780 inverted confocal microscope with a 63x/1.4 numerical aperture oil-immersion objective. <u>Widefield microscopy</u>: fixed, mounted coverslips were imaged at 20X or 40X using a widefield fluorescence microscope (Nikon Eclipse Ti2) with an LED light source. <u>Electron microscopy:</u> resin-embedded ultrathin 70 nm sections on copper grids were imaged on a FEI Tecnai G2 Spirit TEM. Images were acquired in a CCD camera (Eagle).

### 2.12 Recycling measurement by widefield microscopy

HEK293-SNAP-GLP-1R cells seeded on coverslips were treated in serum free medium with agonist (100 nM) or vehicle for 30 minutes, in reverse time order, followed by washing and a variable recycling incubation period. Cells were then labelled using SNAP-Surface-549 (1 µM) in complete medium for 30 minutes on ice to label surface receptor (contributed to by a recycled receptor population and a further population that were not internalised during the agonist stimulation step), fixed with 4% paraformaldehyde, and mounted in Diamond Prolong mounting medium. Slides were imaged by widefield microscopy as above. Five images were taken per coverslip in regions of high confluence using TRITC and FITC filter sets, followed by quantification of surface labelling in Fiji. Cell autofluorescence signal, estimated from a slide with cells that did not undergo SNAP-labelling, was subtracted.

### 2.13 Recycling measurement by confocal microscopy

INS-1 832/3-SNAP-GLP-1R cells on coverslips were treated with agonist (100 nM) or vehicle for 30 minutes in serum-free RPMI, followed by washing. Exendin-4-TMR (100 nM) was then added at the beginning of a 3-hour recycling period, after which cells were fixed with 4% paraformaldehyde, mounted in Diamond Prolong mounting medium, and 3 images acquired per coverslip by confocal microscopy as above. In this assay, total uptake of exendin-4-TMR is indicative of residual surface receptor at the end of exendin-4/lixisenatide pre-treatment (expected to be low) and the cumulative reappearance surface receptor during the recycling period. Quantification was performed in Fiji by expressing the integrated density of specific TMR signal relative to the number of cells for each image.

### 2.14 Lysotracker co-localisation assay

Surface SNAP-GLP-1R was first labelled in INS-1 832/3-SNAP-GLP-1R cells with SNAP-Surface-488 (1 µM) for 30 minutes at 37°C. After washing, cells were treated in serum free RPMI, 11 mM glucose, with agonist or vehicle added in reverse time order. Lysotracker-DND99 (100 nM; Thermo Fisher) was added for the final 30 minutes before fixation and mounting as above. Co-localisation of SNAP-GLP-1R with Lysotracker was quantified from cell-containing image regions as Mander’s coefficient with auto-threshold detection using the Coloc2 algorithm in Fiji. At least 4 images were analysed per coverslip.

### 2.15 Ultrastructural analysis of SNAP-GLP-1R localization by electron microscopy

INS-1 832/3-SNAP-GLP-1R cells on Thermanox coverslips (Agar Scientific) were labeled with 2 μM SNAP-Surface-biotin (a gift from Dr Ivan Corrêa Jr, New England Biolabs), and 5 μg/ml NaN_3_-free Alexa Fluor 488 Streptavidin, 10 nm colloidal gold (Molecular Probes) and stimulated with 100 nM exendin-4 for 1 hour. Conventional EM was performed as previously described (Tomas et al., 2004). Ultrathin 70 nm sections were cut *en face* with a diamond knife (DiATOME) and imaged as above. Electron micrographs were individually thresholded to create binary images displaying only gold particles. All images were then systematically processed to quantify gold-labelled receptors so that multi-particle aggregates would be registered as single large complexes as follows: first, the ImageJ “dilate” algorithm was applied three times so that adjacent gold particles within a complex would coalesce; and second, the ImageJ “particle analysis” algorithm was run on the processed image to quantify the area of all particles and particle aggregates. The number of particles per aggregate was estimated using the determined size of clearly identified single gold particles after image processing. 40 images per treatment were quantified.

### 2.16 Insulin secretion assays

INS-1 832/3 cells were stimulated with agonist for 16-18 hours in complete medium at 11 mM glucose. At the end of the incubation period, a sample of supernatant was removed, diluted, and analysed for secreted insulin by HTRF (Insulin High Range Assay, Cisbio). Insulin secretion was expressed relative to that from cells stimulated with 11 mM glucose alone in the same experiment.

### 2.17 Animal studies

Mice are commonly used to test metabolic effects of GLP-1R agonists (Greig et al., 1999). All animal procedures were approved by British Home Office under the UK Animal (Scientific Procedures) Act 1986 (Project Licence PB7CFFE7A). Lean male C57BL/6 mice (8-16 weeks of age, body weight 25-30 g, purchased from Charles River) were maintained at 21-23°C and light-dark cycles (12:12 hour schedule, lights on at 07:00). *Ad libitum* access to water and normal chow diet (RM1, Special Diet Services) was provided unless otherwise stated. Mice were housed in groups of four, except for prior to food intake assessments when they were individually caged, with one week of acclimatisation prior to experiments.

### 2.18 Intraperitoneal glucose tolerance tests

Mice were lightly fasted (2-3 hours) before the test. Glucose (2 g/kg) was injected via the IP route, either concurrently with agonist or after a specified delay. Blood glucose was recorded at the indicated time-points from the tail vein using a hand-held glucometer (GlucoRx Nexus).

### 2.19 Food intake assay

Individually caged mice were fasted overnight. Diet was returned immediately after IP injection of agonist, and intake monitored by measuring food weight at the indicated time-points.

### 2.20 Pharmacokinetic study

Mice were injected IP with agonist, and a blood sample was subsequently taken from the tail vein into lithium-heparin capillary tubes onto ice. Plasma was separated and stored at −80°C prior to determination of exendin-4 / lixisenatide concentration using an ELISA which equally cross reacts with both peptides (Exendin-4 fluorescent EIA Kit, Phoenix Pharmaceuticals).

### 2.21 Group sizes, randomisation and blinding

*<u>In vitro</u>* <u>experiments</u>: all *in vitro* experiments subjected to statistical analysis were performed with at least *n*=5 biological replicates. Some experiments reported were repeated fewer than 5 times; examples include some corroborative confocal microscopy experiments performed to support quantitative DERET measures of internalisation, optimisation experiments e.g. for BG-SS-Lumi4-Tb, or where the labour-intensive nature of the experiments precluded multiple repeats, such as electron microscopy. Such results should be considered exploratory. Throughout, one biological replicate was considered as the average of technical replicates (2-3) from each assay. Treatments were randomly distributed across microplates to avoid systematic bias. Each treatment to be compared was included in each experiment to allow matched analyses. Due to resource limitations it was not possible to perform *in vitro* treatments in a blinded manner. *<u>In vivo</u>* <u>experiments</u>: all *in vivo* experiments included at least 5 mice per group. Group sizes were determined on the basis of previously established *n* numbers required to demonstrate the size of the effect expected from the *in vitro* results, without formal power calculations. Treatment order was randomly assigned and the average body weight within each group calculated to ensure this did not differ by more than 1 g. The investigator performing the experiment was blinded to treatment allocations.

### 2.22 Data and statistical analysis

The data and statistical analysis comply with the recommendations on experimental design and analysis in pharmacology (Curtis et al., 2018). Quantitative data were analysed using Prism 8.0 (GraphPad Software). Mean ± standard error of mean (SEM), or individual replicates, are displayed throughout. Normalisation was used for some *in vitro* assays to control for unwanted sources of variation between experimental repeats. Specifically, concentration-response data were expressed either relative to the maximal response of a reference compound, or relative to vehicle-treated control; nanoBiT recruitment responses, FRET biosensor responses, and TR-FRET internalisation and recycling kinetic responses were expressed relative to individual well baselines. Concentration-response data were analysed using 4-parameter logistic fits, with basal responses and Hill slopes globally constrained. As exendin-4 and lixisenatide were both full agonists in all signalling assays, comparison of log EC_50_ values (relative potency ratios) was appropriate for determination of signal bias (Kenakin and Christopoulos, 2013); most figures refer to “pEC_50_” values, i.e. modulus of log EC_50_, so that more potent ligands are assigned a higher value. Statistical comparisons were made by t-test or ANOVA as appropriate, either paired or matched depending on the experimental design. Statistical significance was inferred if p<0.05. Image analysis was performed in Fiji as above.

## 3 Results

### 3.1 Lixisenatide displays impaired coupling to intracellular cAMP signalling

The sequences of exendin-4 and lixisenatide are shown in Figure 1A. We used a TR-FRET approach (Emami-Nemini et al., 2013) to measure binding of each ligand to human SNAP-tagged GLP-1R in competition with a fluorescent GLP-1R antagonist, exendin(9-39)-FITC (Supplementary Figure 1A, 1B, 1C). The equilibrium dissociation constants were similar for both agonists (log K_d_ −9.2 ± 0.1 *versus* −9.0 ± 0.1 for exendin-4 and lixisenatide, respectively, p>0.05 by paired t-test, Figure 1B). Calculation of association and dissociation rate constants from kinetic binding experiments in competition with exendin(9-39)-FITC also showed no differences between ligands (Figure 1C). However, in spite of similar binding affinities, cAMP signalling potency for lixisenatide was approximately five times lower than for exendin-4 (log EC_50_ −10.0 ± 0.1 *versus* −9.3 ± 0.1 for exendin-4 and lixisenatide, respectively, p<0.05 by paired t-test, Figure 1D). Potency for GLP-1R endocytosis, measured in parallel with the cAMP signalling experiments using diffusion-enhanced resonance energy transfer (DERET), was also significantly reduced for lixisenatide, but to a more minor degree (log EC_50_ −8.0 *versus* −7.7 ± 0.1 for exendin-4 and lixisenatide, respectively, p<0.05 by paired t-test, see Supplementary Figure 1D for kinetic traces), with analysis of pathway bias confirming that lixisenatide coupling to cAMP was selectively reduced. Confocal imaging confirmed both ligands induced extensive SNAP-GLP-1R endocytosis (Figure 1E).

**Figure 1.**
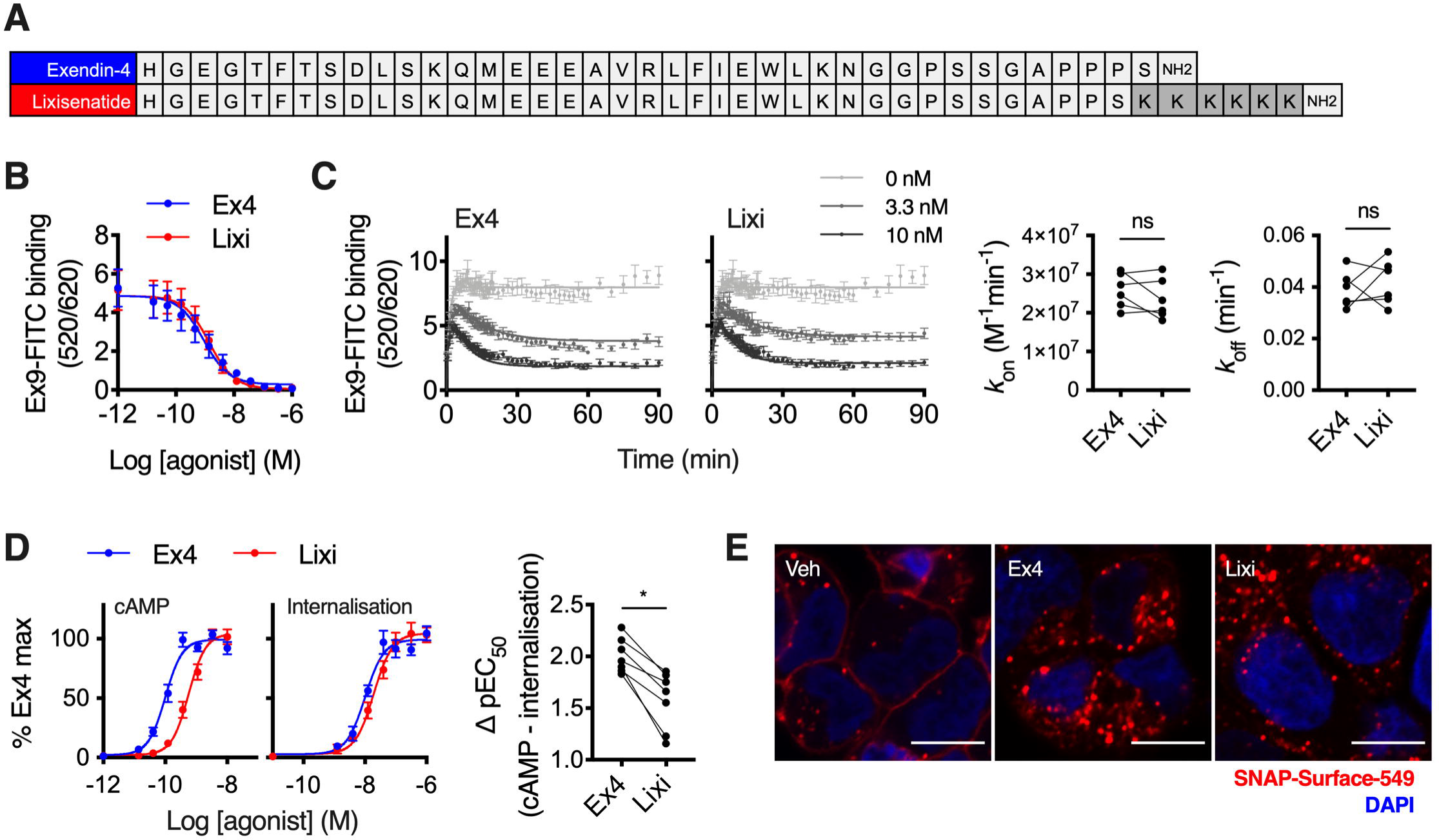
Lixisenatide displays selectively reduced coupling to cAMP signalling. (**A**) Sequences of each ligand in single amino acid code. (**B**) Equilibrium binding experiment using exendin-4 (Ex4) or lixisenatide (Lixi) in competition with 10 nM exendin(9-39)-FITC (Ex9-FITC) in HEK293-SNAP-GLP-1R cells, *n*=5, see also Supplementary Figure 1B. (**C**) Kinetic binding experiment using exendin-4 or lixisenatide in competition with 10 nM exendin(9-39)-FITC in HEK293-SNAP-GLP-1R cells, with calculation of association (*k*_on_) and dissociation (*k*_off_) rate constants, *n*=6, paired t-tests. (**D**) Parallel measurements of cAMP production and GLP-1R internalisation in HEK293-SNAP-GLP-1R cells, 30-minute incubation, *n*=7, 4-parameter fits of pooled data shown, bias analysis shows ΔpEC_50_ for cAMP – internalisation responses, paired t-test. (**E**) SNAP-GLP-1R endocytosis in HEK293-SNAP-GLP-1R cells labelled with SNAP-Surface-549, 30-minute stimulation with 100 nM ligand or vehicle (Veh), size bars: 8 µm, representative confocal images of *n*=2 experiments. *p<0.05 by statistical test indicated in the text. Data shown as mean ± SEM or individual replicates.

As bias can be time point-specific (Klein Herenbrink et al., 2016), we also used the FRET biosensor ^T^Epac^VV^ (Klarenbeek et al., 2011) to obtain real-time readouts of cAMP signalling (Supplementary Figure 1E). Comparison of ^T^Epac^VV^ and DERET potencies at 10-minute intervals indicated the selective loss of cAMP potency with lixisenatide was preserved throughout the stimulation period (Supplementary Figure 1F). Downstream coupling to protein kinase A (PKA) activation was similarly reduced with lixisenatide (Supplementary Figure 1G). These studies indicate that coupling of lixisenatide to cAMP signalling is reduced compared to with exendin-4, in a manner unrelated to receptor occupancy.

### 3.2 Recruitment responses measured using NanoBiT complementation

To attempt to understand why lixisenatide shows reduced cAMP signalling despite similar binding affinity to exendin-4, we also used a NanoBiT complementation approach (Dixon et al., 2016) to measure recruitment of “mini-G proteins” (Wan et al., 2018), and β-arrestin-2, to GLP-1R. We found that both ligands at a supramaximal dose induced robust recruitment of mini-G_s_, with a modestly reduced response with lixisenatide (AUC 292.7 ± 48.0 *versus* 258.0 ± 47.8 for exendin-4 and lixisenatide, respectively, p<0.05 by paired t-test) (Figure 2A). Mini-G_i_ or −G_q_ were not recruited following stimulation with ligand. Recruitment of β-arrestin-2 was also slightly reduced with lixisenatide compared to exendin-4 (Figure 2B; AUC 48.1 ± 13.8 *versus* 41.4 ± 14.6 for exendin-4 and lixisenatide, respectively, p<0.05 by paired t-test). Notably, β-arrestin-2 recruitment signal with both ligands showed more rapid decay than seen with mini-G_s_, which is compatible with a previous report ascribing class A β-arrestin kinetics to the GLP-1R (Al-Sabah et al., 2014). As the ligand concentration used was supramaximal, the implication of these studies is that lixisenatide is a modestly less efficacious ligand than exendin-4 for both G_s_ and β-arrestin-2 recruitment.

**Figure 2.**
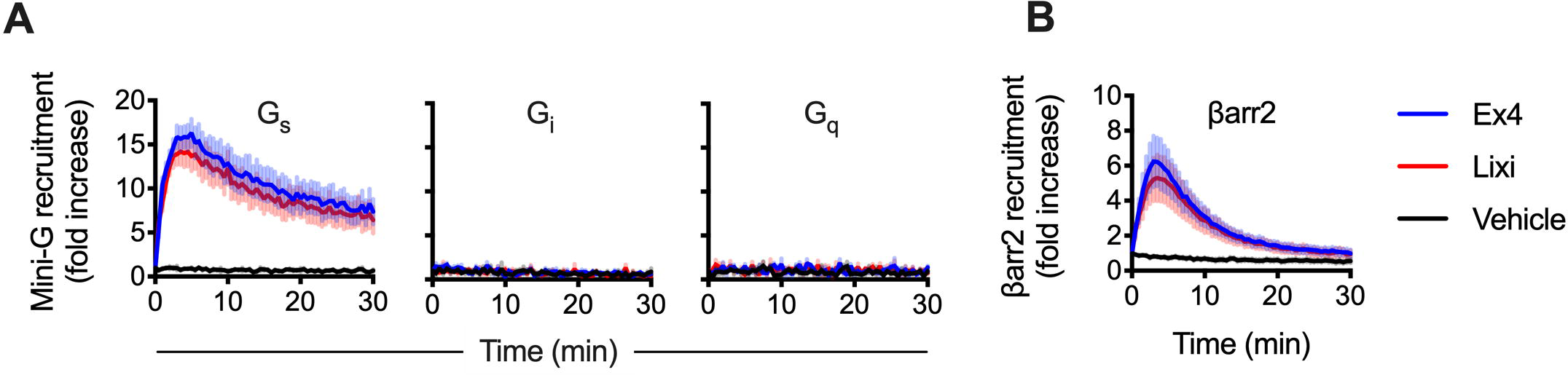
Recruitment of G proteins and β-arrestin-2 to GLP-1R. (**A**) Recruitment of LgBiT-tagged mini-G_s_, −G_i_ and −G_q_ to GLP-1R-SmBit in transiently transfected HEK293T cells treated with 1 µM exendin-4 or lixisenatide, measured by NanoBiT complementation, *n*=5 (mini-G_s_) or *n*=4 (mini-G_i_, −G_q_). (**B**) As for (A) but recruitment of LgBiT-tagged β-arrestin-2 (βarr2), *n*=5. Data shown as mean ± SEM.

### 3.3 Differences in GLP-1R recycling following exendin-4 and lixisenatide treatment

After initial endocytosis, differences in subcellular receptor trafficking can modulate GLP-1R-induced insulin release (Buenaventura et al., 2018; Jones et al., 2018b). GLP-1R is chiefly sorted either towards lysosomal degradation or recycled back to the plasma membrane, with the balance between the two predictive of insulinotropic efficacy of biased agonists (Jones et al., 2018b). To measure GLP-1R recycling, we treated HEK293-SNAP-GLP-1R cells with exendin-4 or lixisenatide, followed by a variable recycling period, after which we applied a SNAP-Surface fluorescent probe to label total surface GLP-1R. This showed that treatment with lixisenatide *versus* exendin-4 was associated with a slower rate of GLP-1R recycling (Figure 3A). To corroborate this finding, we developed a TR-FRET assay to measure SNAP-GLP-1R recycling in real-time using a plate reader (see Figure 3B for a graphical description of the assay principle). The assay uses “BG-SS-Lumi4-Tb”, a cleavable form of the SNAP-labelling TR-FRET donor SNAP-Lumi4-Tb that incorporates a disulfide bond (Supplementary Figure 2A) to allow reversible labelling by release of the fluorophore under mild reducing conditions. The probe performs similarly to non-modified SNAP-Lumi4-Tb for GLP-1R labelling, measurements of fluorescent GLP-1R agonist binding and for internalisation by DERET (Supplementary Figure 2B, 2C, 2D). However, unlike the non-cleavable probe, the cell-impermeant reducing agent Mesna can be used to un-label residual surface receptors, as previously described using non-lanthanide SNAP-probes (Buenaventura et al., 2018), so that fluorescent signal originates exclusively from internal receptors endocytosed during a prior agonist treatment step (see Supplementary Figure 2E, 2F for optimisation of Mesna cleavage). Re-emergence of labelled receptors at the plasma membrane is then detected by TR-FRET using exendin(9-39)-FITC as the acceptor. Using CHO-K1 cells stably expressing SNAP-GLP-1R, which remain adherent during the multiple wash steps, we again found a reduced rate of GLP-1R recycling after lixisenatide compared to exendin-4 treatment (Figure 3C).

**Figure 3.**
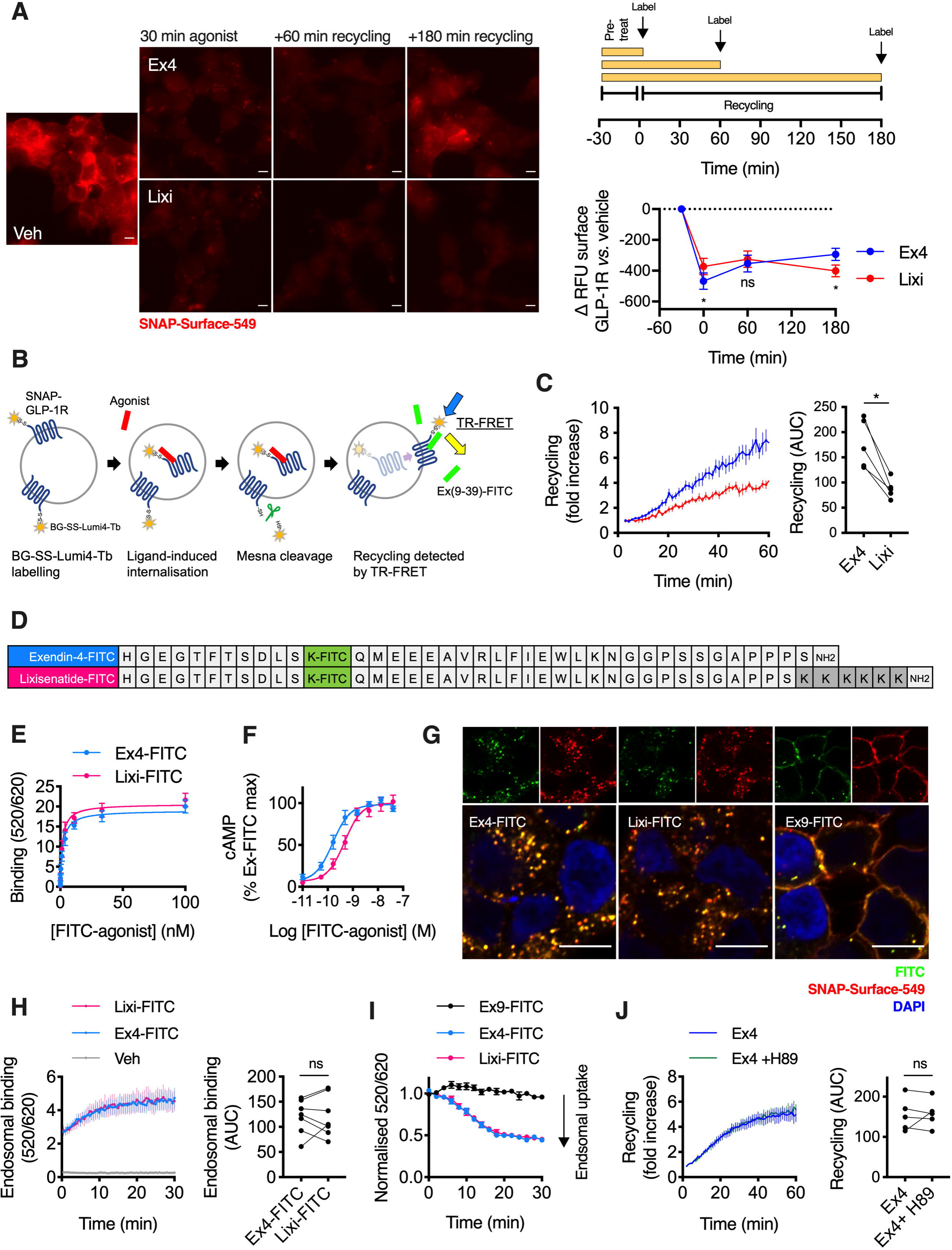
GLP-1R recycling with exendin-4 *versus* lixisenatide. (**A**) Non-labelled HEK293-SNAP-GLP-1R cells were treated with exendin-4 or lixisenatide (100 nM) for 30 minutes, followed by washing, indicated recycling period, and finally labelling of surface GLP-1R with SNAP-Surface-549, representative microscopy images are shown with quantification of recycled receptor as relative fluorescence units (RFU) from *n*=5 experiments, two-way repeat measures ANOVA with Sidak’s test, size bars: 8 µm. (**B**) Principle of TR-FRET recycling assay. (**C**) GLP-1R recycling after exendin-4 or lixisenatide treatment (100 nM, 30 minutes) in CHO-K1-SNAP-GLP-1R cells, measured by TR-FRET, *n*=5, paired t-test for AUC quantification. (**D**) Sequences of each FITC-ligand in single amino acid code. (**E**) Saturation binding of exendin-4-FITC (Ex4-FITC) and lixisenatide-FITC (Lixi-FITC) in HEK293-SNAP-GLP-1R cells, *n*=6. (**F**) cAMP responses in HEK293-SNAP-GLP-1R cells, 30 min incubation, *n*=5, 4-parameter fit shown. (**G**) Confocal microscopy images of HEK293-SNAP-GLP-1R cells labelled with SNAP-Surface-549 and treated with indicated FITC-ligand (100 nM) for 30 minutes, size bars: 8 µm, representative images of *n*=2 independent experiments. (**H**) Endosomal binding of FITC-ligands in HEK293-SNAP-GLP-1R cells, 100 nM ligand treatment for 30 minutes prior to acid wash, *n*=7, paired t-test for AUC quantification. Note that the slight increase in signal over time is likely to represent restoration of “normal” endosomal pH following the prior exposure to pH 2.9. (**I**) TR-FRET signal of FITC-ligands in HEK293-SNAP-GLP-1R cells pre-bound with indicated agonist (100 nM) at 4°C before initiating endocytosis by transfer to 37°C, normalised to initial TR-FRET signal, *n*=5. (**J**) GLP-1R recycling after exendin-4 (100 nM, 30 minutes) treatment in CHO-K1-SNAP-GLP-1R cells with or without prior treatment with H89 (10 µM), measured by TR-FRET, *n*=5, paired t-test for AUC quantification. *p<0.05 by statistical test indicated in the text. Data shown as mean ± SEM or individual replicates.

As pH-dependent dissociation of ligand-receptor complexes within acidic endosomes is known to influence post-endocytic receptor sorting (Borden et al., 1990), we wondered if the impact of low pH might specifically affect interactions made by the positively charged lixisenatide C-terminus, modulating intra-endosomal binding and thereby explaining its different recycling rate. To investigate this possibility, we developed FITC-conjugates of each agonist (Figure 3D) to use as TR-FRET acceptors in intra-endosomal binding assays. Initial assessment showed that the FITC conjugate peptides recapitulated the pharmacological properties of their unmodified counterparts, with similar binding affinity but reduced cAMP signalling potency for lixisenatide-FITC compared to exendin-4-FITC (Figure 3E, 3F). Confocal microscopy demonstrated both exendin-4-FITC and lixisenatide-FITC to be localised within GLP-1R-containing endosomes 30 minutes post-stimulation (Figure 3G); however, in an acid wash assay (Supplementary Figure 2G), TR-FRET signals of each FITC-ligand bound to internalised GLP-1R labelled with SNAP-Lumi4-Tb suggested that endosomal binding of each compound is similar (Figure 3H). Moreover, the progressive loss of TR-FRET signal from agonist pre-bound to surface receptors at low temperature and subsequently endocytosed by return to 37°C was equal for both conjugates (Figure 3I, Supplementary Figure 2H, 2I), also suggesting intra-endosomal dissociation of agonist-receptor complexes does not differ between agonists. Overall, these results suggest the difference in recycling rate between exendin-4 and lixisenatide is not related to differences in persistence of receptor binding within endosomes.

As GPCR recycling can be controlled by PKA (Vistein and Puthenveedu, 2013), we also wondered whether our observation of reduced PKA signalling despite similar occupancy for lixisenatide *versus* exendin-4 (Supplementary Figure 1G) might explain their different recycling rates. However, treatment with the PKA inhibitor H89 did not affect GLP-1R recycling after exendin-4 pre-treatment (Figure 3J).

### 3.4 Divergent effects of exendin-4 and lixisenatide in pancreatic beta cells

Potentiation of glucose-stimulated insulin secretion by beta cells is a major physiological and therapeutic outcome of GLP-1RAs. We used incretin-responsive rat insulinoma-derived INS-1 832/3 beta cells to investigate differences between exendin-4 and lixisenatide in the native cellular context for GLP-1R. As expected, reduced signalling potency for cAMP with lixisenatide was apparent in this cell type (log EC_50_ −9.0 ± 0.1 *versus* −9.5 ± 0.0, p<0.05 by paired t-test, Figure 4A), and also in mouse insulinoma-derived MIN6B1 cells (log EC_50_ −10.5 ± 0.2 *versus* −9.8 ± 0.3, p<0.05 by paired t-test, Figure 4B). Moreover, the pattern of signal bias first demonstrated in HEK293 cells was recapitulated when we expressed SNAP-GLP-1R in INS-1 832/3 cells in which the endogenous GLP-1R was deleted by CRISPR/Cas9 (Naylor et al., 2016), with lixisenatide showing a relative preference for internalisation compared to cAMP signalling (Figure 4C, Supplementary Figure 3A). Similar uptake of exendin-4-FITC and lixisenatide-FITC was observed by confocal microscopy (Figure 4D). To measure GLP-1R recycling in this cell type after pre-treatment with exendin-4 or lixisenatide, we applied fluorescent exendin-4-TMR (Supplementary Figure 3B, 3C) at the beginning of the recycling period following extensive wash of unlabeled agonist. As exendin-4-TMR is rapidly endocytosed by GLP-1Rs that reappear at the cell surface, its intracellular accumulation is indicative of the recycling rate. Again, recycling of GLP-1R after pre-treatment with lixisenatide was less extensive than with exendin-4 (Figure 4E).

**Figure 4.**
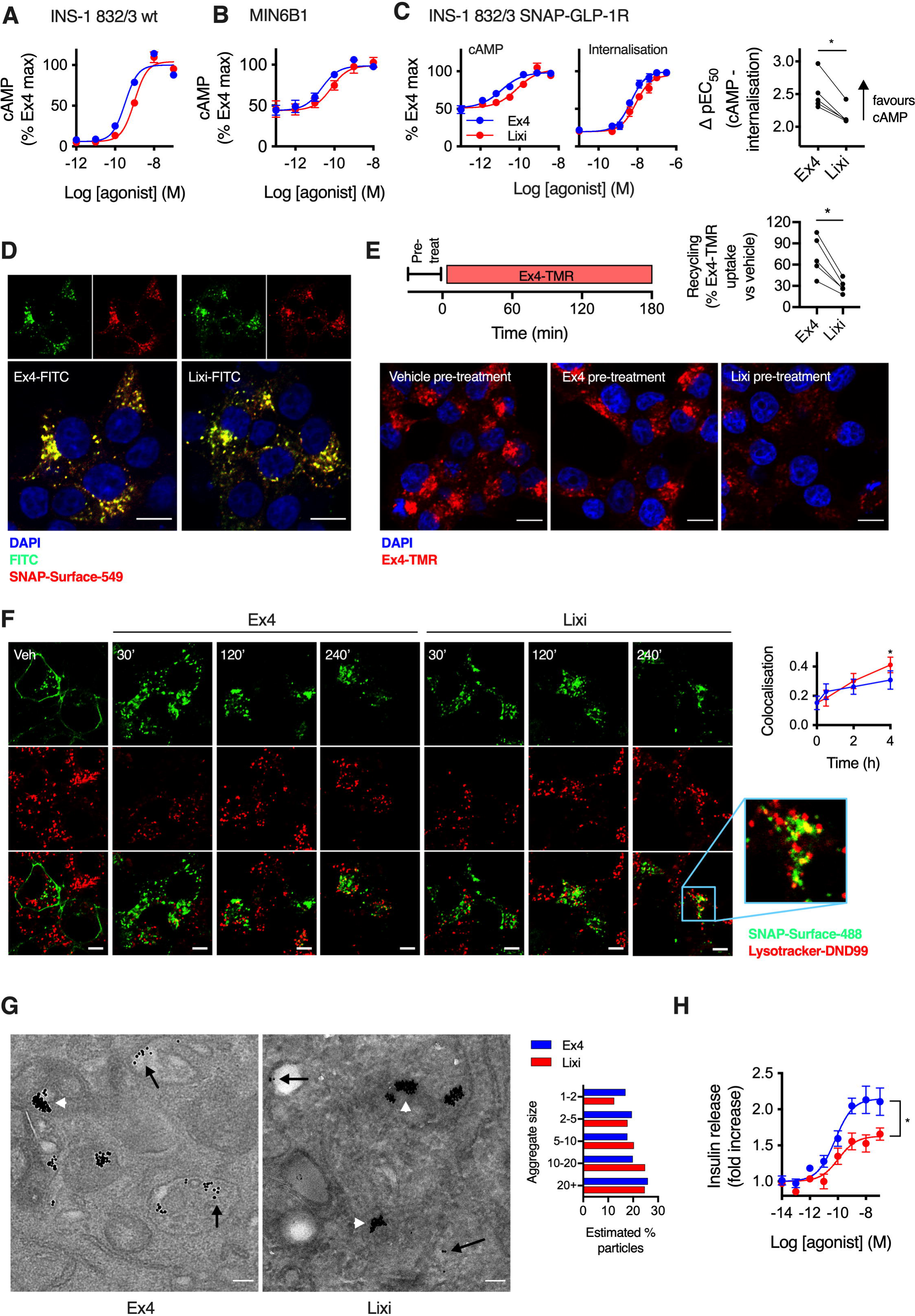
Effects of exendin-4 and lixisenatide in beta cells. (**A**) Acute cAMP responses in wild-type (wt) INS-1 832/3 cells treated with each ligand and 500 µM IBMX for 10 minutes, *n*=6, 4-parameter fits of pooled data shown. (**B**) As for (A) but in MIN6B1 cells and *n*=5. (**C**) Acute cAMP (30 minutes with 500 µM IBMX) and internalisation responses measured by DERET in INS-1 832/3-SNAP-GLP-1R cells, *n*=5, with quantification of bias and comparison by paired t-test, see also Supplementary Figure 3A for DERET traces. (**D**) Confocal microscopy of INS-1 832/3-SNAP-GLP-1R cells labelled with SNAP-Surface-549 and treated with indicated FITC-ligand (100 nM) for 30 min, size bars: 8 µm, representative images of *n*=2 independent experiments. (**E**) GLP-1R recycling measured in INS-1 832/3 SNAP-GLP-1R cells treated with vehicle, exendin-4 or lixisenatide (100 nM, 30 minutes) followed by washout and 3-hour exposure to exendin-4-TMR (Ex4-TMR, 100 nM) during the recycling period, representative images from *n*=5 experiments showing exendin-4-TMR uptake with quantification, paired t-test, size bars: 8 µm. (**F**) Confocal microscopy images of INS-1 832/3-SNAP-GLP-1R cells labelled with SNAP-Surface-488 prior to stimulation with indicated agonist (100 nM), with Lysotracker-DND99 added 15 minutes before the end of the incubation, representative images from *n*=5 experiments with quantification by colocalisation, 2-way repeat measures ANOVA with Sidak’s test, size bars: 5 µm; inset, high magnification area. (**G**) Representative electron micrographs of INS-1 832/3-SNAP-GLP-1R cells labelled prior to stimulation with SNAP-Surface-biotin followed by 10 nm gold-conjugated streptavidin and subsequently treated with exendin-4 or lixisenatide (100 nM, 1 hour), size bars: 0.1 µm; black arrows: single gold particles in endosomes, white arrowheads: lysosomal gold aggregates; quantification of size of aggregates from *n*=40 images per treatment is shown. (**H**) Insulin secretion after 16-hour stimulation with exendin-4 or lixisenatide in INS-1 832/3 cells, *n*=7, 4-parameter fit shown, paired t-test to compare E_max_. *p<0.05 by statistical test indicated in the text. Data shown as mean ± SEM or individual replicates.

Agonist-internalised receptors which do not follow a recycling pathway can be sorted towards lysosomal degradation, and in keeping with this, we found increased colocalisation of SNAP-GLP-1R with a fluorescent lysosomal marker after prolonged treatment with lixisenatide compared to exendin-4 (Figure 4F). Moreover, electron microscopy imaging showed that, following live cell labelling of surface SNAP-GLP-1Rs with a 10 nm gold probe prior to agonist stimulation, gold particles were more likely to be found in larger size aggregates with lixisenatide *versus* exendin-4 treatment (Figure 4G). This pattern of gold aggregation is indicative of probe target lysosomal degradation, as previously demonstrated for the epidermal growth factor receptor (EGFR) (Futter and Hopkins, 1989; Futter et al., 1996). As excessive loss of surface GLP-1Rs without compensatory increases in recycling can limit insulinotropic efficacy (Jones et al., 2018b), we measured cumulative insulin secretion after overnight treatment with each agonist (Figure 4H). Consistent with this paradigm, maximal insulin release with lixisenatide was reduced. Thus, the distinct pharmacological properties of lixisenatide and exendin-4 translate to functional differences in beta cells.

### 3.5 Lixisenatide is less effective *in vivo*

GLP-1RAs are primarily used for the treatment of T2D to reduce glycaemia and promote weight loss through appetite reduction. We assessed the glucoregulatory effects of each ligand at varying doses in mice via IP glucose tolerance tests (IPGTTs) performed immediately and 6 hours after agonist treatment, to identify acute and delayed effects. We found that the anti-hyperglycaemic effect of exendin-4 was greater than equimolar lixisenatide (Figure 5A, 5B, 5C). Measurements of food intake in overnight-fasted mice also showed that the anorectic effect of lixisenatide is reduced compared to exendin-4 (Figure 5D, 5E, 5F). As expected from published data (Kolterman et al., 2005; Distiller and Ruus, 2008), these differences did not appear attributable to pharmacokinetics, as plasma concentrations of each agonist measured at 6 hours after IP injection were similar (Figure 5G).

**Figure 5.**
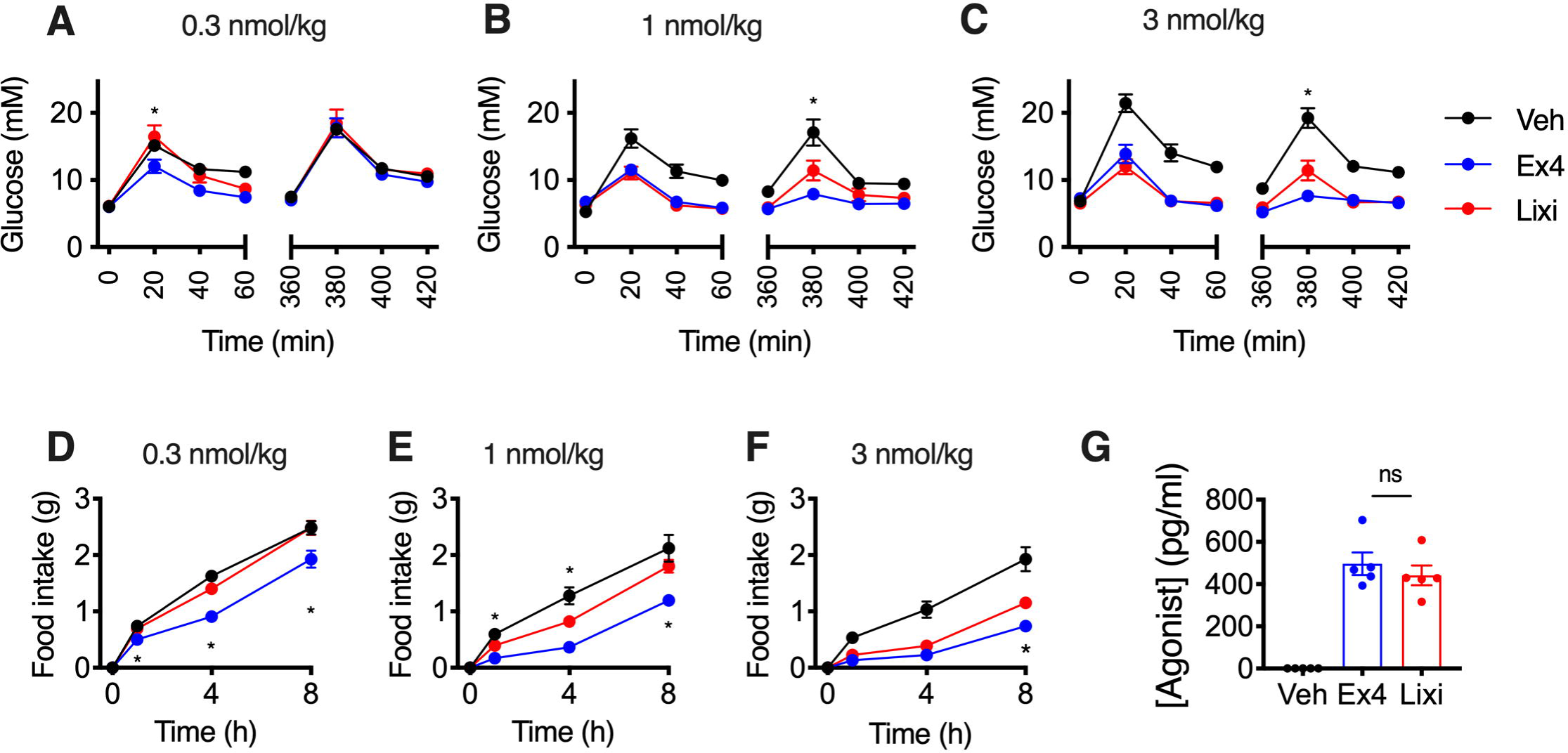
*In vivo* effects of exendin-4 *versus* lixisenatide. (**A**) IP glucose tolerance tests performed in lean mice at immediate and delayed (6-hour) time-points after IP administration of 0.3 nmol/kg ligand or vehicle (saline), 2 g/kg glucose, *n*=8/group, two-way repeat measures ANOVA with Tukey’s test showing comparison between exendin-4 and lixisenatide at each time-point. (**B**) As for (A) but with agonist dose of 1 nmol/kg. (**C**) As for (A) but with agonist dose of 3 nmol/kg and *n*=12/group. (**D**) Cumulative food intake in overnight-fasted lean mice after IP administration of 0.3 nmol/kg ligand or vehicle (saline), *n*=8/group, two-way repeat measures ANOVA with Tukey’s test showing comparison between exendin-4 and lixisenatide at each time-point. (**E**) As for (D) but with agonist dose of 1 nmol/kg. (**F**) As for (D) but with agonist dose of 3 nmol/kg. (**G**) Agonist plasma concentrations in mice at 6 h after IP administration of indicated agonist (100 nmol/kg) or vehicle (saline), *n*=5/group, unpaired t-test. *p<0.05 by statistical test indicated in the text. Data shown as mean ± SEM or individual replicates.

### 3.6 Biased agonists of lixisenatide display similar characteristics to their exendin-4-derived counterparts

Based on the observation that single amino acid substitutions close to the exendin-4 N-terminus can influence agonist-induced GLP-1R signalling and trafficking behaviours (Jones et al., 2018b), we generated “lixi-phe1” and “lixi-asp3” (Supplementary Figure 4A). In the context of exendin-4, the -phe1 substitution is known to reduce binding affinity, reduce GLP-1R endocytosis, accelerate GLP-1R recycling, enhance insulin release, and improve anti-hyperglycaemic efficacy, while exendin-asp3 shows opposing characteristics (Jones et al., 2018b). Here, we found that, as for exendin-phe1, lixi-phe1 is a lower affinity ligand than its -asp3 counterpart (Figure 6A, Supplementary Figure 4B). Similarly, acute cAMP signalling potency was reduced for both -phe1 variants (Figure 6B). However, when the reduced affinity was accounted for, exendin-phe1 and, to a lesser extent, lixi-phe1, were more efficiently coupled to cAMP production than the -asp3 versions (Figure 6C). Striking differences in GLP-1R trafficking were noted, with robust and equal endocytosis induced by exendin-asp3 and lixi-asp3 when measured by DERET, but very little from the exendin-phe1 or lixi-phe1 (Figure 6D, Supplementary Figure 4C; some GLP-1R internalisation with -phe1 ligands was detectable by confocal microscopy [Figure 6E]). Recycling of endocytosed receptor was rapid with both -phe1 analogues (lixi-phe1 less so than exendin-ph1) and slow with the -asp3 equivalents (Figure 6F). The recycling rate correlated well with binding affinity when plotted on a log-log scale (Figure 6G). In INS-1 832/3 cells, insulin release after a 16 hour incubation at a maximal agonist concentration was notably higher with both lower affinity -phe1 *versus* corresponding -asp3 ligands (Figure 6H), contrasting with the greater acute signalling potency with the latter in the same cell line (Supplementary Figure 4D). Correspondingly, both -phe1 analogues exerted superior anti-hyperglycaemic effects in an IPGTT performed 8 hours after agonist administration, with exendin-phe1 being the most effective (Figure 6I). The -asp3 peptides showed no effect in comparison to vehicle. In keeping with our previous observation that the trafficking effects of biased GLP-1R agonists tend to exert a more pronounced effect on glucose regulation than on food intake (Jones et al., 2018b), anorectic effects of -phe1 and -asp3 peptides were similar (Figure 6J), although there was a non-significant trend towards reduced efficacy for both -phe1 compounds at early timepoints which was reversed at 8 hours.

**Figure 6.**
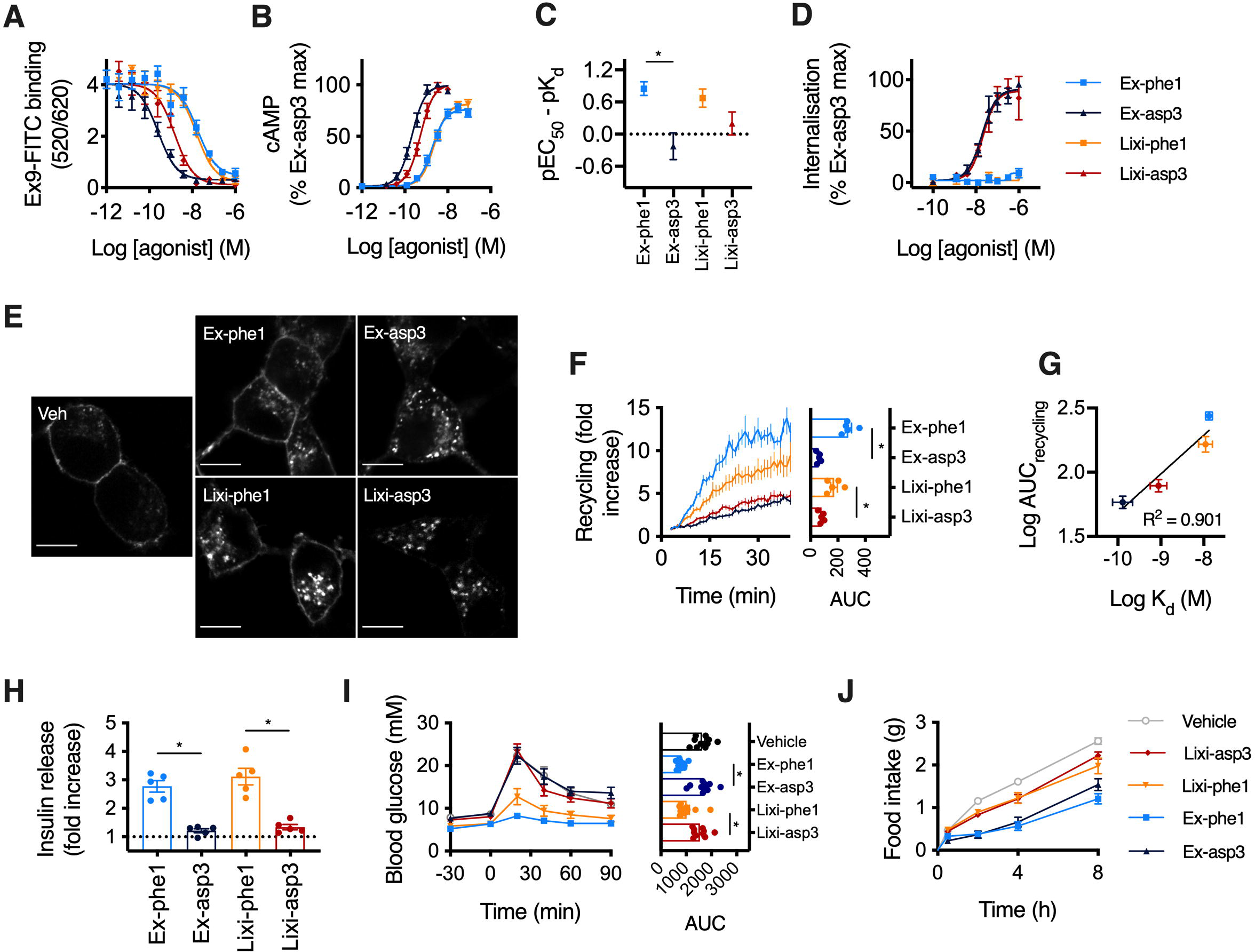
Comparison of biased exendin-4 and lixisenatide analogues. (**A**) Equilibrium binding experiment using unlabeled agonists in competition with 10 nM exendin(9-39)-FITC in HEK293-SNAP-GLP-1R cells, *n*=5, see also Supplementary Figure 4B. (**B**) cAMP responses in HEK293-SNAP-GLP-1R cells, 30-minute incubation, *n*=5, 4-parameter fits of pooled data shown. (**C**) Quantification of coupling of receptor occupancy to cAMP signalling for each ligand, quantified by subtracting pEC_50_ from data shown in (B) from pK_d_ shown in (A), with error propagation, one-way ANOVA with Sidak’s test. (**D**) GLP-1R endocytosis in HEK293-SNAP-GLP-1R cells measured by DERET, 30-minute incubation, *n*=5, 4-parameter fits shown, see also Supplementary Figure 4C. (**E**) Confocal microscopy images showing SNAP-GLP-1R endocytosis in INS-1 832/3-SNAP-GLP-1R cells labelled with SNAP-Surface-549, 100 nM agonist, 30-minute incubation, size bars: 8 µm, representative images from *n*=2 experiments. (**F**) GLP-1R recycling in CHO-K1-SNAP-GLP-1R cells, measured by TR-FRET, 30-minute pre-treatment with 100 nM agonist followed by Mesna cleavage of surface BG-SS-Lumi4-Tb and monitoring of recycling by TR-FRET, *n*=5, AUC compared by one-way randomized block ANOVA with Tukey’s test. (**G**) Comparison of ligand affinity with recycling rate using data from (A) and (F), linear regression performed to determine goodness of fit. (**H**) Insulin secretion in INS-1 832/3 cells treated at 11 mM glucose ± indicated agonist (100 nM) for 16 hours, expressed relative to response without agonist, *n*=5, one-way randomised block ANOVA with Tukey’s test. (**I**) IP glucose tolerance tests performed in lean mice at delayed (8-hour) time-points after IP administration of 2.4 nmol/kg ligand or vehicle (saline), 2 g/kg glucose, *n*=10 (vehicle, exendin-phe1, lixi-asp3) or 11 (exendin-asp3, lixi-phe1) per group, one-way ANOVA with Tukey’s test comparing AUC for each treatment. (**J**) Cumulative food intake in overnight-fasted lean mice injected IP with 2.4 nmol/kg ligand or vehicle (saline), *n*=12/group. *p<0.05 by statistical test indicated in the text. Data shown as mean ± SEM or individual replicates.

Two main conclusions can be made from this set of experiments: 1) sequence substitutions close to the N-termini of both parent peptides consistently result in a linked set of *in vitro* and *in vivo* agonist characteristics with the potential to improve their therapeutic properties, and 2) this effect is most marked for exendin-4, for which the differences between -phe1 and -asp3 analogues was generally greater than for lixisenatide.

## 4 Discussion

In this study we performed a side-by-side pharmacological evaluation of two closely related GLP-1RAs, both of which are in routine clinical usage and differ structurally only by the hexalysine C-terminal extension in lixisenatide. We found that, despite similar binding affinity, coupling of lixisenatide to cAMP signalling was reduced compared to exendin-4. After similar levels of GLP-1R endocytosis induced by both peptides, recycling of GLP-1R was slower after lixisenatide treatment, with apparent preferential targeting to a degradative lysosomal pathway. This resulted in a reduction in insulinotropic efficacy and the ability to control blood glucose 6-8 hours after dosing in mice. Therefore, the structural differences between the C-termini of each peptide appear to have some functional importance.

Our finding here of reduced cAMP signalling potency with lixisenatide matches our earlier observation (Jones et al., 2018b). We also found reduced cAMP potency in rat INS-1 832/3 and mouse MIN6B1 cells, arguing against a species-specific effect. Unfortunately, in the absence of a high level of confidence of the position of the exendin-4 C-terminus in published GLP-1R structural studies (Liang et al., 2018), a robust molecular explanation for the signalling deficit of lixisenatide remains elusive. Runge *et al*. found the exendin-4 Trp-cage does not directly interact with the GLP-1R ECD (Runge et al., 2008), although analysis of photo-crosslinking data using the full length receptor showed that the flexible exendin-4 C-terminus might snake over the top of the ECD and form interactions with Phe80, Tyr101 and Phe103 (Koole et al., 2017). Ligand-specific C-termini could plausibly alter this interaction, and it is possible to envisage resultant changes to the orientation of the ligand N-termini, with knock-on effects on interactions with the receptor core required for full activation. In an alternative interpretation of the aforementioned photo-crosslinking data (Koole et al., 2017), the Trp-cage acts *in trans* across GLP-1R homodimers, which might lead to changes in its oligomeric state. As formation of GPCR oligomers can influence coupling to signalling intermediates (Milligan et al., 2019), including for GLP-1R (Harikumar et al., 2012; Buenaventura et al., 2019), this provides a hypothetical mechanism by which the two ligands could signal differently despite similar occupancy. Using NanoBiT complementation, we found subtle differences in GLP-1R coupling to both G_s_ and β-arrestin-2 between exendin-4 and lixisenatide, although it is not clear whether this is sufficient to explain the significant differences in potency for cAMP signalling. Of note, we found that neither ligand was capable of inducing detectable recruitment of G_q_, raising questions about the importance of signalling via this G protein in GLP-1R responses (Shigeto et al., 2015), although we cannot exclude cell type-specific effects.

The other key finding here pertains to the post-endocytic trafficking of the two ligands, which was evaluated across different cell systems using a variety of complementary approaches. Despite similar internalisation profiles, GLP-1R recycling after lixisenatide treatment was notably slower in both HEK293 and INS-1 832/3 beta cells. In contrast, a higher degree of lixisenatide-stimulated GLP-1Rs tended to progressively co-localise post-endocytosis with the lysosomotropic fluorescent probe Lysotracker, indicating preferential targeting of the receptor towards a degradative pathway. This was in agreement with the increased level of intracellular gold-conjugated SNAP-tag probe aggregation detected by electron microscopy, suggesting enhanced tendency for lysosomal degradation of the lixisenatide-stimulated receptor. These recycling phenotypes partly recapitulate the differences previously observed with biased exendin-4-derived GLP-1R agonists (Jones et al., 2018b), which were linked to diminished insulin secretion efficacy. Indeed, we found in the present study that maximal insulin secretion was reduced when beta cells were exposed to lixisenatide *versus* exendin-4 over a sustained exposure period. However, despite these broadly similar sets of findings, the mechanism for slowing of GLP-1R recycling with lixisenatide did not appear to depend on greater binding affinity, a factor that was previously demonstrated to influence the recycling of biased exendin-4-derived GLP-1R agonists (Jones et al., 2018b). We specifically aimed to address the possibility of protonation of the hexalysine scaffold of lixisenatide in acidic conditions leading to altered intra-endosomal agonist dissociation, but could find no evidence for this. Moreover, PKA signalling, previously implicated in the control of β_2_-adrenergic receptor recycling (Vistein and Puthenveedu, 2013) and indeed linked to downregulation of GLP-1R in beta cells under hyperglycaemic conditions (Rajan et al., 2015), differed between exendin-4 and lixisenatide, but this did not appear to underpin their recycling phenotypes. We did not examine posttranslational modifications linked to target degradation such as ubiquitination, but note that the GLP-1R is not ubiquitinated by treatment with exendin-4 (Jones et al., 2018a). The reason for the increased lysosomal post-endocytic targeting of the GLP-1R with lixisenatide therefore remains unclear.

Our observations of generally reduced biological effect of lixisenatide for physiologically important readouts suggest that these pharmacological differences are indeed translated to differences in downstream responses. In particular, we found reduced efficacy for sustained insulin secretion with lixisenatide using an *in vitro* beta cell system, as well as reduced anti-hyperglycaemic and anorectic effects in mice. As the “advantages” of exendin-4 *in vivo* were detectable acutely, in a dose-dependent manner, it is likely that they are partly attributable to agonist potency differences rather than the post-endocytic trafficking phenotypes. However, the potential for GLP-1R endocytosis to influence pharmacodynamics of exendin-4 has previously been modelled (Gao and Jusko, 2012), and whilst this model focussed on receptor internalisation, differences in recycling and degradation rates could plausibly be linked via similar mechanisms.

Differentiation in the therapeutic profiles of GLP-1RAs in humans has been noted on many occasions (Aroda, 2018). A head-to-head comparison in patients with T2D showed numerically greater HbA1c reduction and weight loss with exenatide compared to lixisenatide (Rosenstock et al., 2013). Exenatide showed a beneficial effect on cardiovascular outcomes, albeit with borderline significance (Holman et al., 2017), whereas a separate trial of lixisenatide found no evidence of benefit (Pfeffer et al., 2015). However, understanding the link between the receptor pharmacology that we observed in our study and real-world performance of each agonist is hampered by the different dosing and administration schedules (10 µg twice daily for exenatide, or weekly as a sustained release preparation, compared to 20 µg once daily for lixisenatide).

Following the distinctive effects we previously observed with biased GLP-1RAs derived from exendin-4 (Jones et al., 2018b), we developed biased lixisenatide-derived compounds based on a similar design. The -phe1 substitution in both exendin and lixisenatide configurations displayed favourable characteristics including reduced internalisation and fast recycling, which translated to improved insulin secretion *in vitro* and significantly better anti-hyperglycaemic effect *in vivo*. These observations add to the evidence that changes to GLP-1RA N-termini are capable of inducing significant signal bias (Zhang et al., 2015; Jones et al., 2018b; Fremaux et al., 2019). The data presented herein highlight however how changes at additional regions within the ligand can further control the impact of these N-terminal substitutions, with the effect on GLP-1R recycling being less dramatic with lixi-phe1 than with exendin-phe1, and with corresponding differences in their anti-hyperglycaemic performance in mice. As for the base exendin-4 and lixisenatide ligands, the reason for these differences may relate to how the ligand C-termini affect the orientation of the N-terminus relative to the core of the receptor, but structural studies will be required to establish if this is the case.

In summary, our study provides insights into specific signalling and trafficking differences of two GLP-1RAs in routine clinical use, linking these characteristics to their effects *in vivo*. The precise molecular mechanisms underpinning these differences remains to be elucidated.

## Acknowledgements

This work was funded by an MRC project grant to B.J., A.T., S.R.B. and G.A.R. The Section of Endocrinology and Investigative Medicine is funded by grants from the MRC, BBSRC, NIHR, an Integrative Mammalian Biology (IMB) Capacity Building Award, an FP7-HEALTH-2009-241592 EuroCHIP grant and is supported by the NIHR Biomedical Research Centre Funding Scheme. The views expressed are those of the author(s) and not necessarily those of the funder. B.J. was also supported by the Academy of Medical Sciences, Society for Endocrinology and an EPSRC capital award. D.J.H. was supported by a Diabetes UK R.D. Lawrence (12/0004431) Fellowship, a Wellcome Trust Institutional Support Award, MRC Confidence in Concept, MRC (MR/N00275X/1 and MR/S025618/1) Project and Diabetes UK (17/0005681) Project Grants. This project has received funding from the European Research Council (ERC) under the European Union’s Horizon 2020 research and innovation programme (Starting Grant 715884 to D.J.H.).

