## Supplementary Information for "Differences in signalling, trafficking and glucoregulatory properties of glucagon-like peptide-1 receptor agonists exendin-4 and lixisenatide"

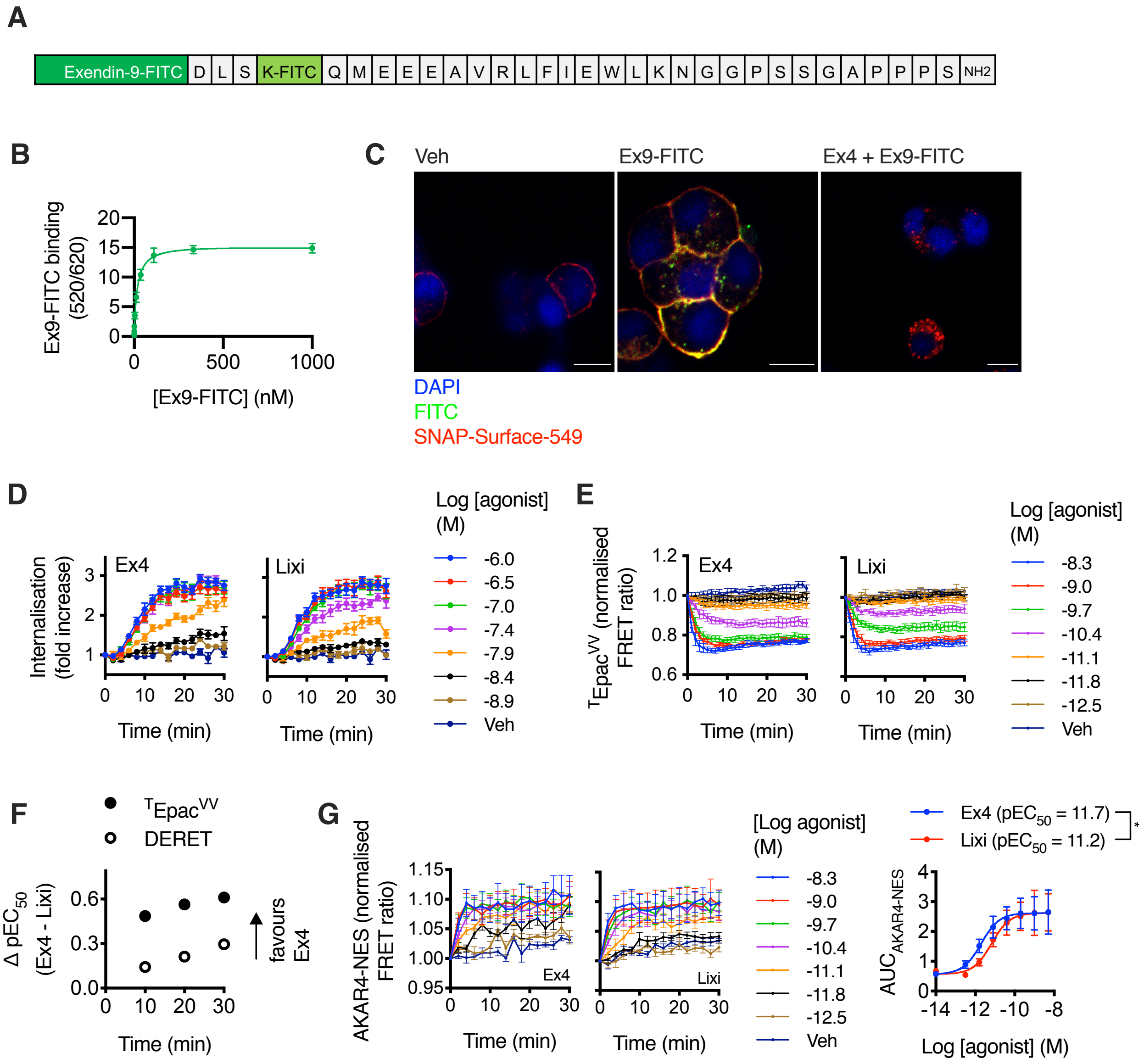


**Supplementary Figure 1.** (**A**) Sequence of exendin(9-39)-FITC in single amino acid code. (**B**) Saturation binding of exendin(9-39)-FITC in HEK293-SNAP-GLP-1R cells, *n*=5, performed in parallel with experiments shown in Figure 1B. (**C**) Confocal microscopy images showing specific exendin(9-39)-FITC binding (100 nM, 30 minutes) to SNAP-Surface-549-labelled INS-1 832/3 SNAP-GLP-1R cells with or without prior treatment with exendin-4 (10 μM). (**D**) DERET internalisation traces with indicated concentration of agonist in HEK293-SNAP-GLP-1R cells, *n*=7, relates to Figure 1D. (**E**) cAMP responses measured using $^{T}Epac^{VV}$ biosensor in HEK293-SNAP-GLP-1R cells with indicated concentration of agonist, normalized to individual well baselines, *n*=5. (**F**) Analysis of bias at 3 timepoints from (D) and (E), with dose responses constructed from averaged data across all experimental repeats split into 10-min bins to derive single values for $pEC_{50}$, lixisenatide responses were subtracted from those of exendin-4 at each time-point. (**G**) Cytoplasmic PKA signalling in CHO-K1-SNAP-GLP-1R cells stably expressing AKAR4-NES biosensor stimulated with indicated concentration of agonist, *n*=5, 4-parameter fit of AUC shown with $pEC_{50}$ values compared by paired t-test. *$p<0.05$ by statistical test indicated in the text. Data indicated as mean ± SEM.

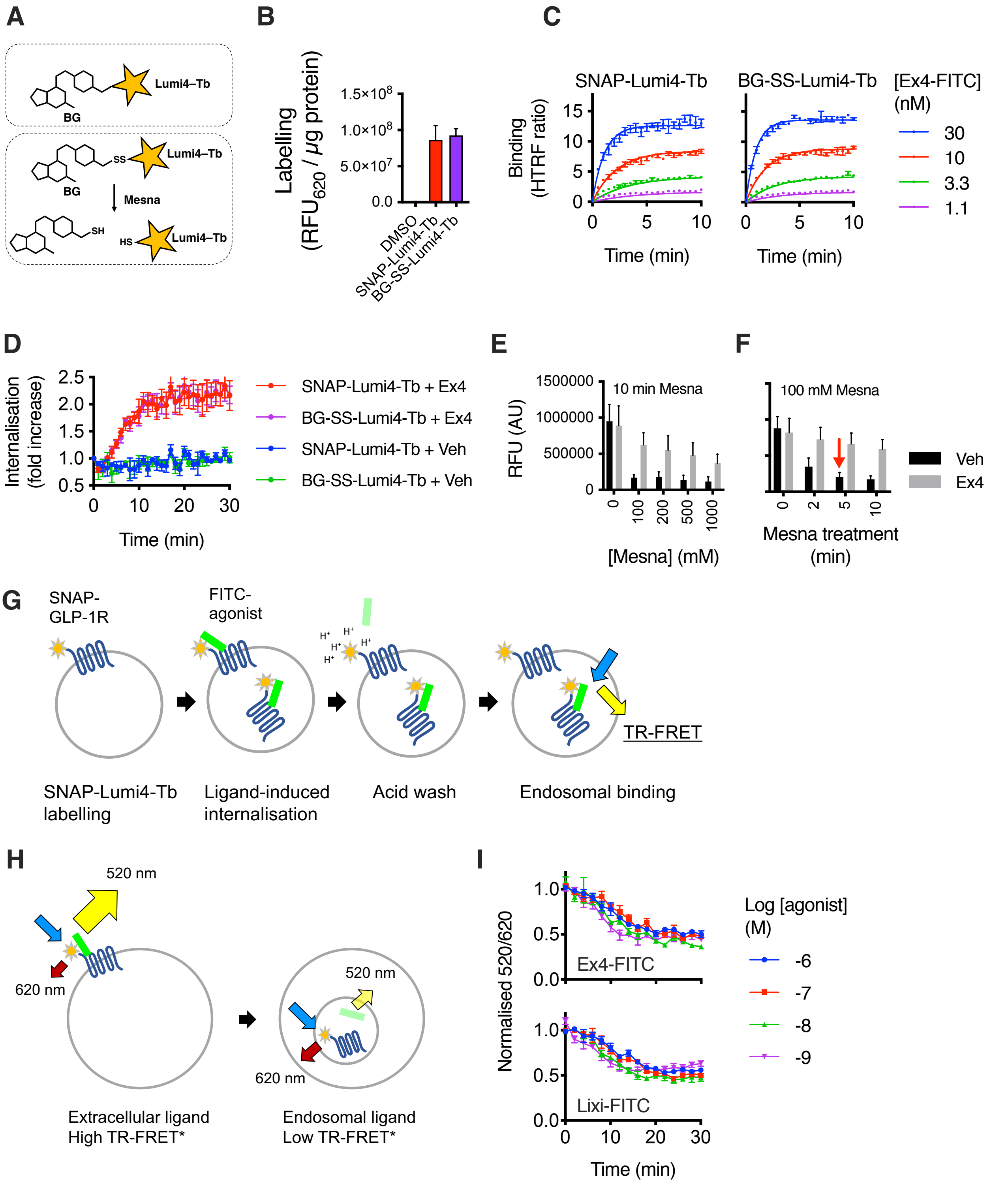


**Supplementary Figure 2.** (**A**) Cartoon depicting cleavage of BG-SS-Lumi4-Tb using Mesna. (**B**) Labelling efficiency of SNAP-Lumi4-Tb and BG-SS-Lumi4-Tb (both 40 nM) in HEK293-SNAP-GLP-1R cells, normalised to protein content, *n*=2. (**C**) Kinetic binding of exendin-4-FITC in HEK293-SNAP-

GLP-1R cells labelled with SNAP-Lumi4-Tb or BG-SS-Lumi4-Tb, *n*=3. (**D**) Exendin-4-induced GLP-1R internalisation measured by DERET in HEK293-SNAP-GLP-1R cells labelled with SNAP-Lumi4-Tb or BG-SS-Lumi4-Tb, *n*=3. (**E**) Optimisation of Mesna cleavage of surface SNAP-GLP-1R in CHO-K1-SNAP-GLP-1R cells labelled with BG-SS-Lumi4-Tb and treated ± exendin-4 (100 nM, 30 minutes) before treatment with Mesna at indicated concentration for 10 minutes at 4°C, *n*=2. (**F**) As for (E) but with fixed Mesna dose for indicated incubation period; red arrow highlights how maximum cleavage was achieved by 5 minutes. (**G**) Principle of endosomal binding experiment shown in Figure 3H. (**H**) Principle of FITC-ligand uptake experiment shown in Figure 3I. (**I**) FITC-ligand uptake in HEK293-SNAP-GLP-1R cells pre-bound with indicated ligand and dose at 4°C before triggering endocytosis by transfer to 37°C, *n*=3. Data indicated as mean ± SEM.

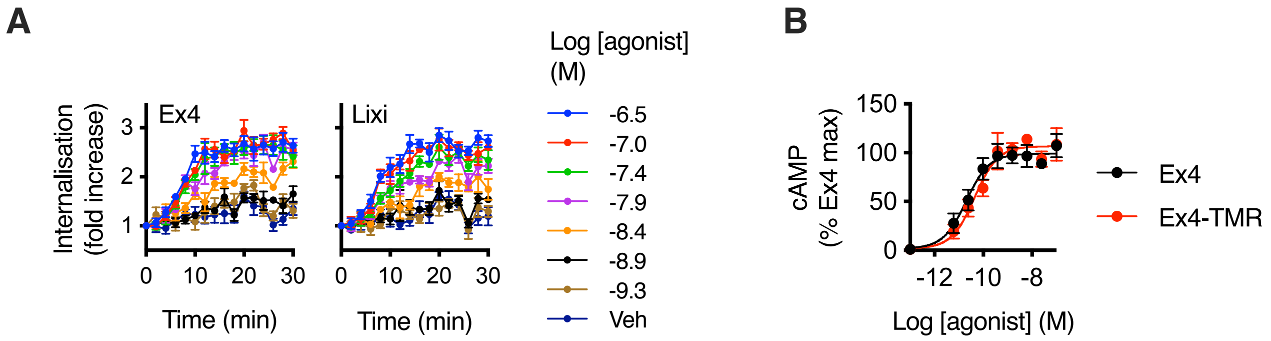


**C**

| | | | | | | | | | | | | | | | | | | | | | | | | | | | | | | | | | | | | | | | | |
|---|---|---|---|---|---|---|---|---|---|---|---|---|---|---|---|---|---|---|---|---|---|---|---|---|---|---|---|---|---|---|---|---|---|---|---|---|---|---|---|---|
| Exendin-4 | H | G | E | G | T | F | T | S | D | L | S | K | Q | M | E | E | E | A | V | R | L | F | I | E | W | L | K | N | G | G | P | S | S | G | A | P | P | P | S | NH2 |
| Exendin-4-TMR | H | G | E | G | T | F | T | S | D | L | S | K-TMR | Q | M | E | E | E | A | V | R | L | F | I | E | W | L | K | N | G | G | P | S | S | G | A | P | P | S | S | NH2 |

**Supplementary Figure 3.** (**A**) DERET internalisation traces with indicated concentration of agonist in INS-1 832/3-SNAP-GLP-1R cells, *n*=5, relates to Figure 4C. (**B**) cAMP responses of exendin-4 and exendin-4-TMR in HEK293-SNAP-GLP-1R cells, 30 min stimulation, 4-parameter fits of pooled data shown, *n*=4. (**C**) Amino acid sequence of exendin-4 and exendin-4-TMR in single letter amino acid code.

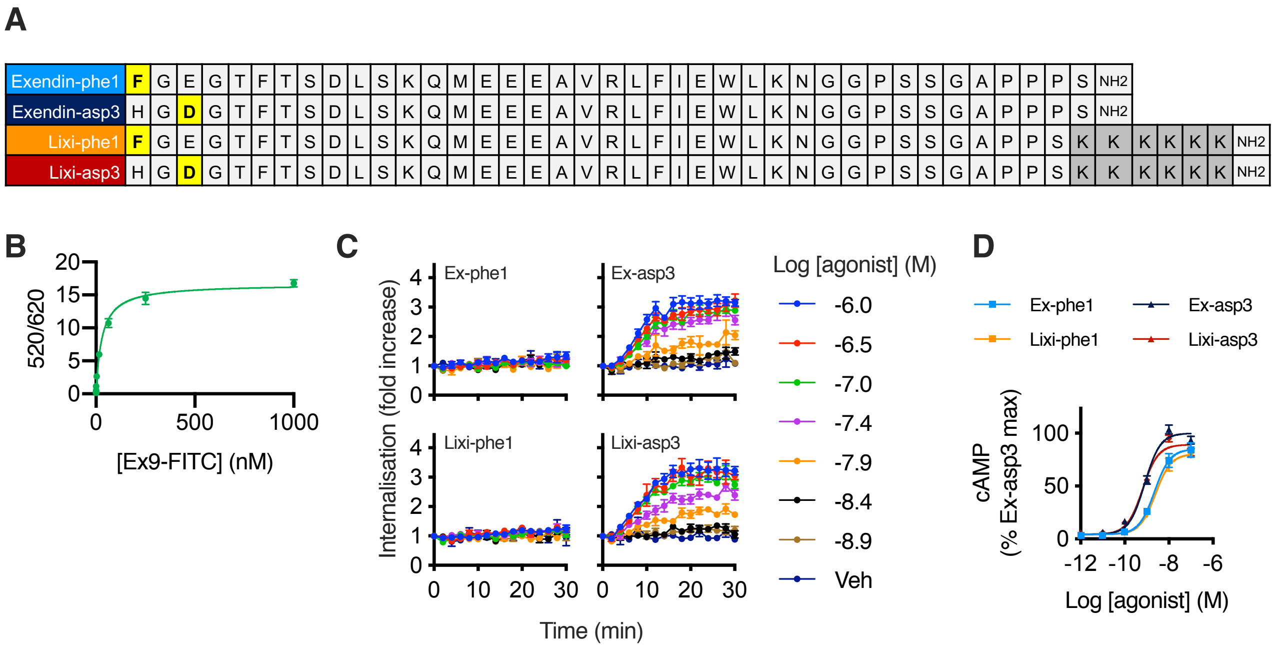


**Supplementary Figure 4.** (**A**) Peptide sequences of biased exendin-4 and lixisenatide analogues in single letter amino acid code. (**B**) Saturation binding of exendin(9-39)-FITC in HEK293-SNAP-GLP-1R cells, *n*=5, performed in parallel with experiment shown in Figure 6A. (**C**) DERET internalisation traces with indicated concentration of agonist in HEK293-SNAP-GLP-1R cells, *n*=5, relates to Figure 6D. (**D**) Acute cAMP responses in INS-1 832/3 cells treated with each ligand and 500 μM IBMX for 10 minutes, normalised to exendin-asp3 response, *n*=4, 4-parameter fits of pooled data shown.
